# Mechanism-guided quantification of LINE-1 reveals p53 regulation of both retrotransposition and transcription

**DOI:** 10.1101/2023.05.11.539471

**Authors:** Alexander Solovyov, Julie M. Behr, David Hoyos, Eric Banks, Alexander W. Drong, Jimmy Z. Zhong, Enrique Garcia-Rivera, Wilson McKerrow, Chong Chu, Dennis M. Zaller, Menachem Fromer, Benjamin D. Greenbaum

## Abstract

Somatic activity of LINE-1 (L1) mobile elements has been implicated in cancer etiology, which may be related to the loss of p53-mediated regulation as a result of *TP53* mutations. Quantifying the mechanisms of L1 regulation in cancer has been challenging. Here, we build a statistical model of L1 regulation by simultaneously quantifying L1 retrotransposition, L1 expression, and the fitness costs of mutated *TP53* with precision. We first developed Total ReCall, an algorithm specifically tailored to the mechanisms of L1 reintegration, to detect L1 insertions from short-read whole-genome sequencing. Applying Total ReCall to high-quality data consisting of >750 paired tumor and normal samples from The Cancer Genome Atlas (TCGA) shows high L1 insertion heterogeneity among tumor types, with increased retrotransposition burden in lung squamous cell carcinoma, head and neck, and colon cancers. We next assessed the active RNA expression of intact L1 in >9,000 TCGA tumor samples, establishing, for the first time, a clear correlation between L1 expression and retrotransposition. Finally, we integrated the number of L1 insertions, L1 expression and a mathematical model of *TP53* fitness into a multi-modal model of p53- mediated mechanisms of L1 regulation. We show that *TP53* mutations enable retrotransposition both by disinhibiting L1 expression and enabling its reintegration and quantify the relative weights of this dual regulatory role. We demonstrate how mechanism-based multi-modal modeling applied at scale can statistically disentangle the complex interplay between canonical driver events in tumor evolution and retrotransposon activity.

## Introduction

More than half of the human genome is composed of repeat sequences^1–3^. Normally, various processes such as epigenetic repression silence these repeats^4^ but oncogenesis is associated with disruptions to these pathways^5, 6^. In cancer, repeats can be re-expressed as RNA, translated in some cases into protein, and may be actively involved in genome instability and cancer immunogenicity^7–12^. The LINE-1 (L1) element is an especially interesting class of repeats which possesses the ability to reinsert itself via retrotransposition at new loci in the human genome^13^. An intact L1 element is ∼6 kb in length, but most of the >1 million copies that comprise ∼17% of the human genome are 5’ truncated^2^ .There are just over 100 L1s capable of coding for the full length ORF1p and ORF2p proteins, the latter functioning as the reverse transcriptase and endonuclease^1^ needed for L1 activity.

Despite its frequent reintegration in cancer genomes^11^, measuring L1 retrotransposition from short-read sequencing data to accurately quantify its movement in cancer poses technical challenges due to the many genomic copies of highly similar L1 sequences, as well as different modes of the retrotransposition process (Fig. 1a-d) making it difficult to disambiguate the source of L1-containing sequencing reads. While methods have been developed for detecting somatic retrotransposition events^11, 14, 15^ we set out to build a first-principles approach that accounts for the specific retrotransposition cycle of L1 (Fig. 1). The resulting “Total ReCall” method relies on two key signals that arise when aligning sequencing reads to the reference genome (Fig. 1g): (i) reads that span the retrotransposon insertion site breakpoint will result in chimeric reads which contain portions of both the insertion site sequence from the human genome as well as the inserted L1 sequence, and lead to “soft clipped” alignments, and (ii) paired-end reads that arise from fragments spanning the inserted L1 sequence will often have one read map near the insertion site, but the other read map to one of the many L1 sequences elsewhere in the genome, even to a different chromosome or a distant site on the same chromosome, resulting in “discordant read pairs”.

**Figure 1.**
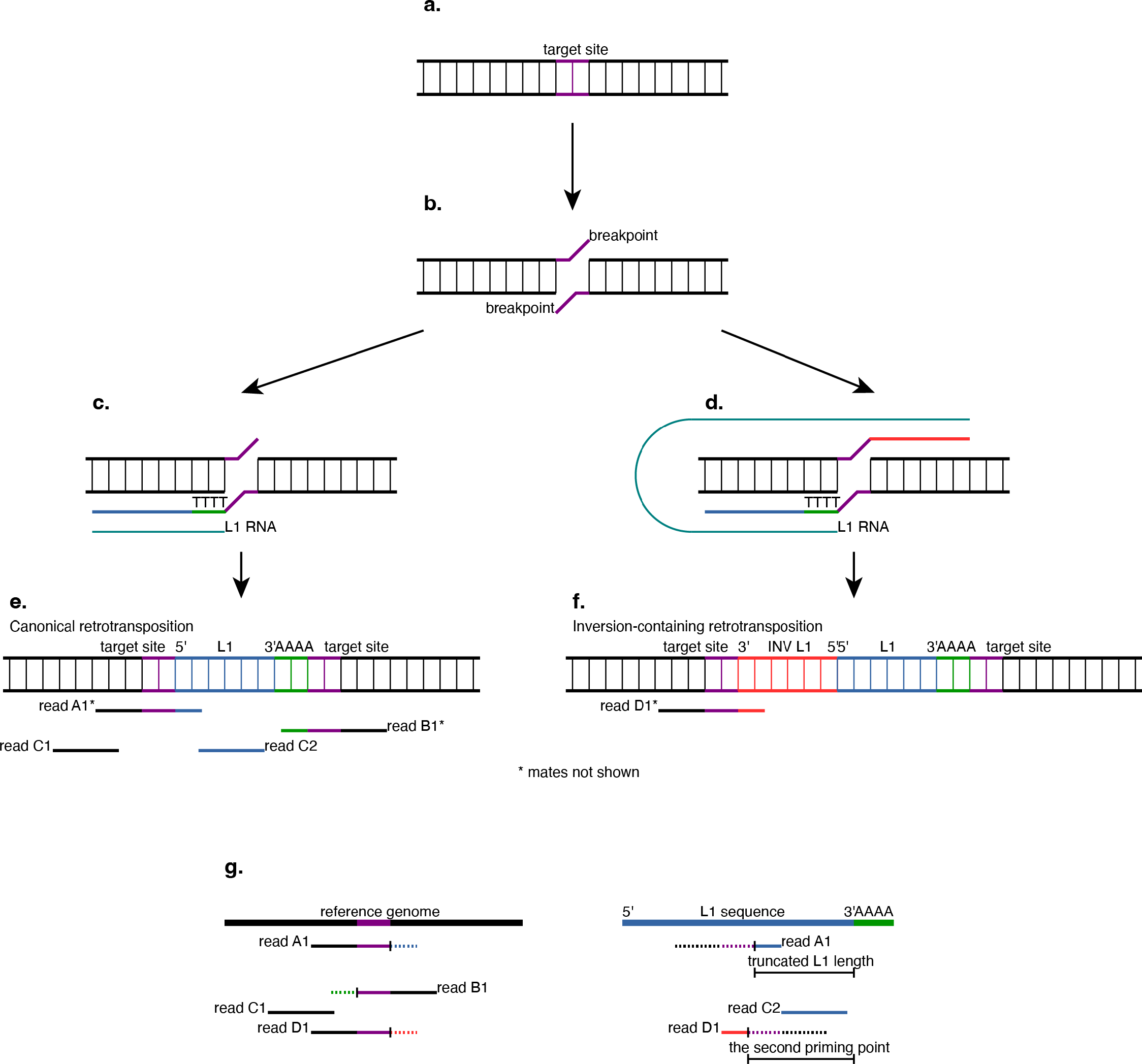
Schematic of insertion of a non-LTR retrotransposon. a) Haploid copy of the genome before retrotransposition. Purple, target site for future insertion. b) Endonuclease breaks each strand of DNA, typically at two nearby but distinct positions. The locus between the two breakpoints is the “target site”. The relative location of the two breakpoints in this figure will result in a target site duplication. c) L1 RNA is reverse transcribed directly into the genome resulting in the synthesis of the single-stranded cDNA starting from the 3’ end of the L1 transcript and extending a variable length towards the 5’ end of the L1 transcript. Teal, L1 RNA. Green, reverse transcribed poly(T) cDNA. Blue, reverse transcribed L1 cDNA. d) In some cases, double priming occurs resulting in the simultaneous reverse transcription of different parts of L1 transcript into the two strands of the genome. Red, reverse transcribed L1 cDNA on the opposite gDNA strand. e-f) Genome after synthesis of the second strand of DNA and repair. The original synthesis strands of cDNA are always complementary to the L1 transcript. Components of the L1 sequence are annotated with respect to the top strand of the genome. Purple, target site which is duplicated following repair. Red, newly inserted L1 sequence that was synthesized on the top strand and is therefore reverse complemented with respect to L1 RNA. Blue, newly inserted L1 sequence that was synthesized on the bottom strand. Green, newly inserted poly(A). Paired-end reads originating from the modified genomes are shown. e) The process shown in panel (c) results in a (possibly truncated at the 5’ end) “canonical” retrotransposition. f) The double priming process shown in panel (d) results in an “inversion-containing” retrotransposition. The resulting genomic sequence has two L1 fragments in opposite orientations. g) Mapping of reads A-D to the unmodified (reference) genome lacking the transposon insertion (left) and the transposon sequence (right). Left, tails of reads A1, B1, and D1 that come from the novel transposon are clipped (shown as dashed lines). Right, read C2 and the clipped tails of reads A1 and D1 align to the transposon sequence. The clipped tail of read B1 contains only poly(T). In the absence of inversion (as in panel (e), captured by read A1), the alignment between the clipped sequence and the transposon sequence reflects the length of the newly inserted transposon. When inversion occurs (as in panel (f), captured by read D1), such an alignment will only reflect the position where the second priming occured.

We applied Total ReCall to assess somatic L1 retrotransposition prevalence in a pan-cancer cohort of 765 paired tumor-normal samples across 22 tissue types from The Cancer Genome Atlas (TCGA), assessing heterogeneity across types. To achieve a clear picture of the L1 life cycle in cancer, we aimed to accurately quantify L1 RNA expression simultaneously with retrotransposition, since RNA not only encodes the protein machinery for retrotransposition but also acts as the substrate for new genomic copies of L1. To do so, we used the L1EM mechanistic model of L1-driven transcription (“active expression”) at intact L1 copies in the genome^16^. Using this, we assessed heterogeneity of active expression of L1 RNA in 9,011 tumor samples across 32 tissue types, and we found similar patterns between the RNA levels of L1 and the abundance of its retrotransposition. *TP53*, the most frequently mutated gene in cancers^17^, encodes a protein that functions to maintain genome stability, including suppression of L1 retrotransposition^18^. Studies involving genetic manipulation of *TP53* in human cell lines and animal models have demonstrated that loss of p53 function leads to derepression of retroelements as seen with common *TP53* cancer mutations^19, 20^. We integrated combined L1 insertional and transcriptional data with *TP53* mutational status and fitness^21^, which allowed us to model this critical regulatory mechanism at large scale. We find that p53 plays a dual role in restraining retrotransposition, both by repression of L1 transcription and by regulation of its integration into the genome.

## Results

### Total ReCall uses clipped sequences to accurately identify somatic L1 insertions

We designed the Total ReCall algorithm (Fig. 1) to analyze patterns in paired-end whole-genome sequencing (WGS) data to detect “canonical” retrotransposition events (where a single stretch of L1 starting from the 3’ end is inserted in the genome; Fig. 1e), as well as “inversion-containing” retrotranspositions (a genomic insertion that contains part of an L1 sequence adjacent to an inverted part of L1; Fig. 1f; see Methods). To derive somatic insertions in tumors, the algorithm looks for read patterns (Fig. 1g) that are specific to the tumor sample and not found in normal tissue from the same patient. For validation, we tested Total ReCall using a child-mother-father trio sequenced with both Illumina short-read and PacBio long-read technologies as part of the NIST-led Genome in a Bottle (GIAB) Consortium^22^, using the latter as ground truth for retrotransposition events (see Methods). We found that Total ReCall on the short-read data had high sensitivity (71%, Extended Data Fig. 1a-b) for detection of long-read-based trio retrotransposition calls, and the inferred lengths of the retrotransposed sequence insertions (estimated for canonical events only) were highly correlated (Pearson correlation R > 0.99, p < 1x10^-10^, Extended Data Fig. 1c).

Applying Total ReCall to TCGA, we found a total of 4,357 tumor-specific somatic retrotranspositions in our quality-controlled (see Methods) dataset of 765 tumors (mean of 5.7 calls per sample; Fig. 2a). As a negative control, we looked for insertions found in the normal tissue but not present in the matched tumor, which yielded only 100 such instances (mean of 0.13 calls per sample), consistent with a low number of false positives. Approximately two thirds (65.5%) of tumor samples were found to have no somatic retrotransposition calls (in the negative control, 91.6% of normal samples had no normal-specific calls). Because many tumor samples were sequenced at higher depths than their paired normal samples, we next confirmed that the tumor samples had a significantly higher rate of somatic L1 calls than the normal samples even when adjusting for sequencing depth in various ways (Extended Data Fig. 2).

**Figure 2.**
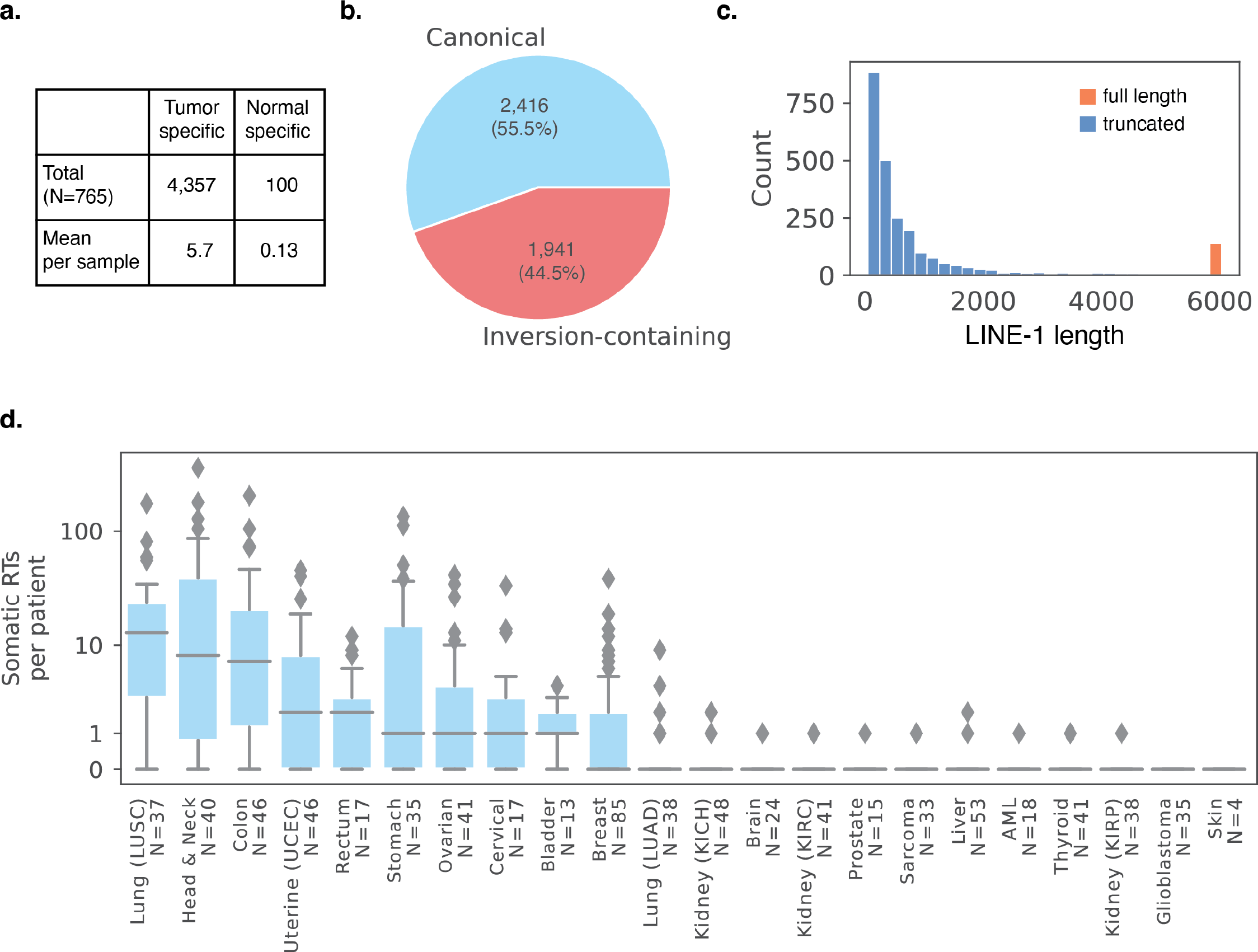
L1 retrotransposition calls from Total ReCall throughout TCGA. N = 765 tumor normal pairs. a) Total number of retrotranspositions identified across 765 tumor and paired normal samples. b) Breakdown of total canonical or inversion-containing insertions identified within the tumor-specific somatic retrotranspositions. c) Estimated length of inserted L1 within the canonical, tumor-specific retrotranspositions. Blue, truncated insertions. Orange, full-length insertions. d) Somatic tumor-specific L1 retrotranspositions in each sample (“RT burden”) grouped by tumor type. Center line indicates median. Blue box indicates interquartile range. Points more than 1.5 x IQR away from the blue box are shown as individual outliers. Tumor types are sorted in descending order by median somatic RT burden.

Among the somatic retrotranspositions detected, slightly over half (2,416 of 4,357, 55.5%) had a canonical orientation of inserted L1 sequence that is consistent with single-stranded priming (Fig. 2b, Fig. 1c), while the remaining (1,941, 44.5%) showed evidence of inversion-containing integrations that result from double-stranded priming (Fig. 1d). This inversion-containing frequency is higher than previous estimates of approximately 19-25% within germline L1 polymorphisms^15, 23^, possibly reflecting the higher levels of genomic instability in tumors. As expected, the length of inserted L1 sequence (estimated from the 2,416 canonical insertions, Fig. 1g) shows a pattern of 5’ decay, with over 94% containing a 5’-truncated L1 sequence, and only 6% approaching full length of a reverse-transcribed L1 transcript (defined here as having at least 97% of the L1Hs consensus [5,833 of 6,032 bases], Fig. 2c). We thus estimate the “mortality rate” for the L1 life cycle in cancer (i.e., the fraction of new L1 integrations that fail to insert the full L1 sequence) as being nearly 19 out of 20.

### Pan-cancer survey reveals highest L1 retrotransposition “burden” in lung squamous cell carcinoma

In our analysis of 765 tumors across the 22 tissue types available in the quality-controlled TCGA data, we found a high degree of heterogeneity in the number of somatic retrotranspositions per sample, dubbed the “RT burden”. In particular, lung squamous cell carcinoma had the highest median RT burden of 13 (Fig. 2d). The next highest, head and neck squamous cell carcinoma tumors had a median RT burden of 8, and the highest mean burden (32.6 RTs / sample) of any tumor type in the dataset. After these, colon adenocarcinoma samples had a median RT burden of 7, and uterine corpus endometrial carcinoma and rectal adenocarcinoma samples both had a median of 2. Stomach, ovarian, cervical, and bladder tumors all had a median RT burden of 1, while breast cancers had a median of 0, but 44% of samples did have at least one somatic RT. In contrast, for the remaining tumor types we did not detect any somatic retrotranspositions in 364 of 388 samples.

Note, however, that without sufficient quality control (QC, see Methods) of the whole-genome sequencing data, the patterns we observed were obscured. In particular, our QC-pipeline removed tumor-normal pairs for which sequencing had been performed earlier in the TCGA project, when read lengths were shorter and thus would not allow the detection of clipped reads (the critical component of Total ReCall), giving the appearance of 0 retrotranspositions in these samples (see Methods). Unfortunately, this filtering removed all esophageal carcinoma and uveal melanoma data from our analysis, cases which are likely to have many RT events^11, 14^.

### L1 RNA expression also differs by tumor types but is related to RT burden

Throughout TCGA, high-quality tumor RNA-seq data is available from 9,011 distinct individuals across 32 tumor types, and normal RNA-seq from 719 individuals across 23 tumor types (see Methods). We quantified locus-level L1 RNA “active” expression (transcription driven by the L1 promoter) from these 9,730 samples using L1EM tool^16^ and aggregated expression levels (transcripts per million, TPM) across fully intact loci (as curated by L1Base^24^) to calculate the total relative abundance of intact L1 RNA present in each sample.

We found that lung squamous cell carcinoma has the greatest median gain of L1 expression in tumor samples with respect to paired normal (tumor median 33.0 TPM, normal median 2.5 TPM, p < 1x10^-10^ one-sided Mann-Whitney U test), while esophageal samples have the highest expression of L1 RNA overall among both tumor and normal samples (tumor median 41.9 TPM, normal median 11.1 TPM, p = 5x10^-5^ one-sided Mann-Whitney U test, Fig. 3a). Prostate tissue had the next highest levels of L1 RNA in normal samples after esophagus, and no significant gain of L1 RNA in the prostate adenocarcinoma samples (tumor median 10.0 TPM, normal median 9.5 TPM, p = 0.12 one-sided Mann-Whitney U test).

**Figure 3.**
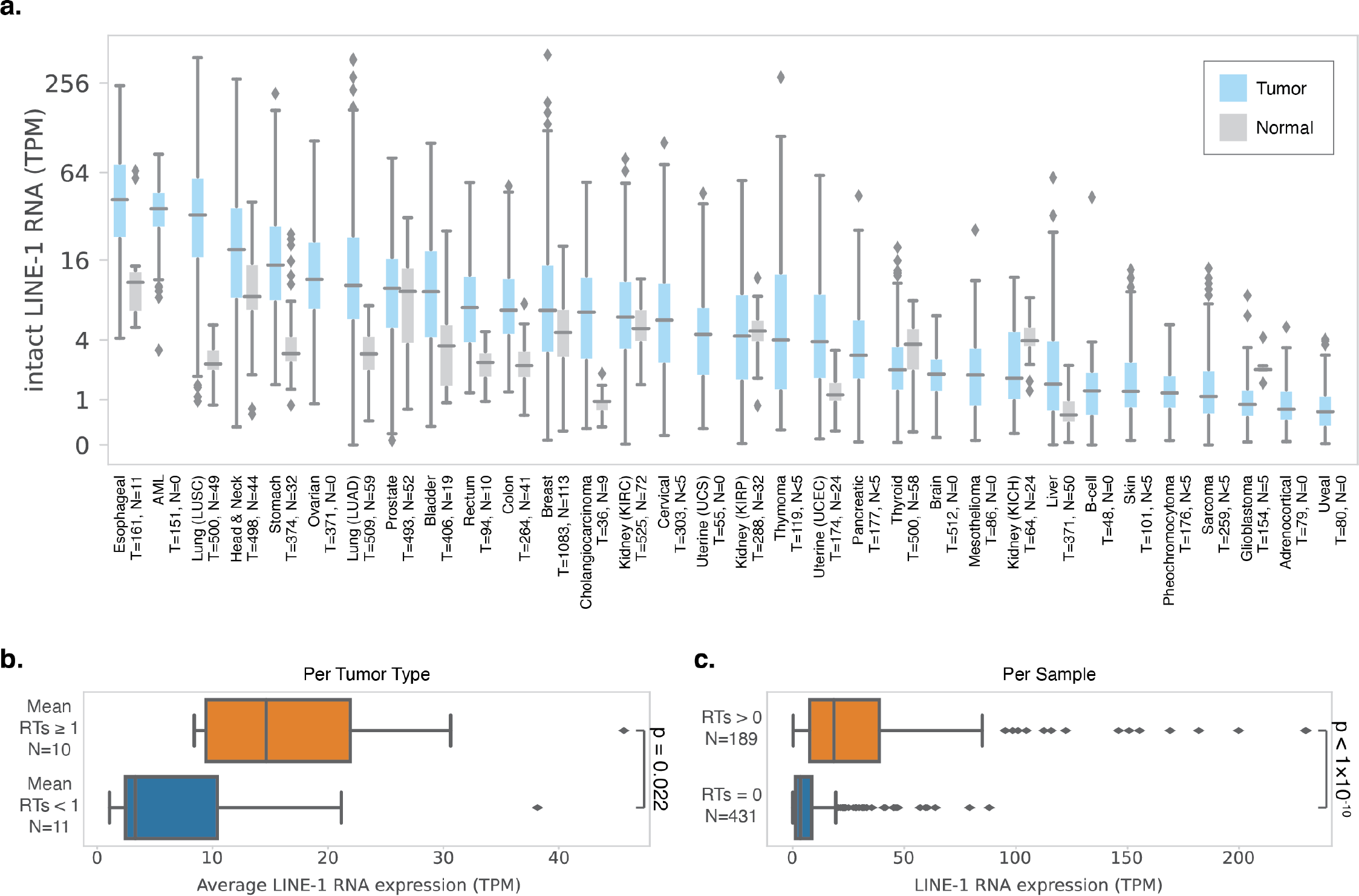
L1 RNA expression from L1EM throughout TCGA. a) Estimated expression of intact L1 RNA in each sample by tumor type. Data not shown for any tumor and sample type with fewer than 5 samples. Total N = 9,715. Blue, tumor samples. Gray, normal samples. Tumor types are sorted in descending order by median tumor-level expression. b) Average expression of intact L1 RNA per tumor type grouped by average retrotransposition count per tumor type. Orange, tumor types with an average of at least one retrotransposition per sample. Blue, tumor types with an average of less than one retrotransposition per sample. c) Expression of intact L1 RNA grouped by retrotransposition count per sample. Orange, samples with any retrotransposition calls. Blue, samples with no retrotransposition calls. a-c, Center line indicates median. Blue, gray, or orange box indicates interquartile range. Points more than 1.5 x IQR away from the IQR box are shown as individual outliers. b-c, P calculated from two-sided Mann-Whitney U test.

Next, to relate L1 retrotransposition to L1 expression, we start by aggregating signals within tumor types and comparing summary statistics. For the 21 tumor types with at least 5 tumor samples in both the WGS and RNA-seq datasets, we ranked each tumor type twice, first based on average L1 RNA expression, then by average RT burden. We found that tumor types with higher retrotransposition ranks tended to also have higher L1 RNA rank (Pearson correlation coefficient R = 0.57, Extended Data Fig. 3a). The 10 tumor types with an average RT burden of at least 1 per sample have higher median L1 RNA expression (median of 14.8 TPM), while the 11 tumor types with an average below one retrotransposition per sample have lower L1 RNA expression (median of 3.3 TPM, p = 0.022, two-sided Mann-Whitney U test, Fig. 3b). We next compared the subset of 620 cancer patients where both retrotransposition and expression could be assessed, i.e., both WGS and RNA-seq had been performed. At the individual sample level, tumors with at least one retrotransposition had significantly higher L1 RNA expression (median 18.0 TPM, N=189) than those with none (median 3.5 TPM, N=431; p < 1x10^-10^, two-sided Mann-Whitney U test, Fig. 3c).

### Pan-cancer statistical model finds that p53 limits retrotransposition by repression of LINE- 1 transcription and regulation of integration

To incorporate the orthogonal signal of *TP53* mutational status into our analyses, we first adjusted our estimates for L1 RNA and RT burden per sample based on sequencing quality metrics to minimize technical biases that may make distinguishing biological relationships more difficult (see Methods, Extended Data Fig. 3b-c). Consistent with the relationship between L1 RNA and L1 RT in Fig. 3b-c, the adjusted values for these also significantly correlate with each other for the tumors with both WGS and RNA-seq data (N = 620, Pearson correlation R = 0.55, p < 1x10^-10^, Extended Data Fig. 3d). We then used the *TP53* alteration status as defined by cBioPortal^25, 26^ to group our WGS and RNA-seq datasets into p53 mutant and wild type categories. The p53 mutant groups exhibited significantly higher L1 RNA and L1 RT burden than the wild type samples (p < 1 x10^-10^ for both, two-sided Mann-Whitney U test, Fig. 4).

**Figure 4.**
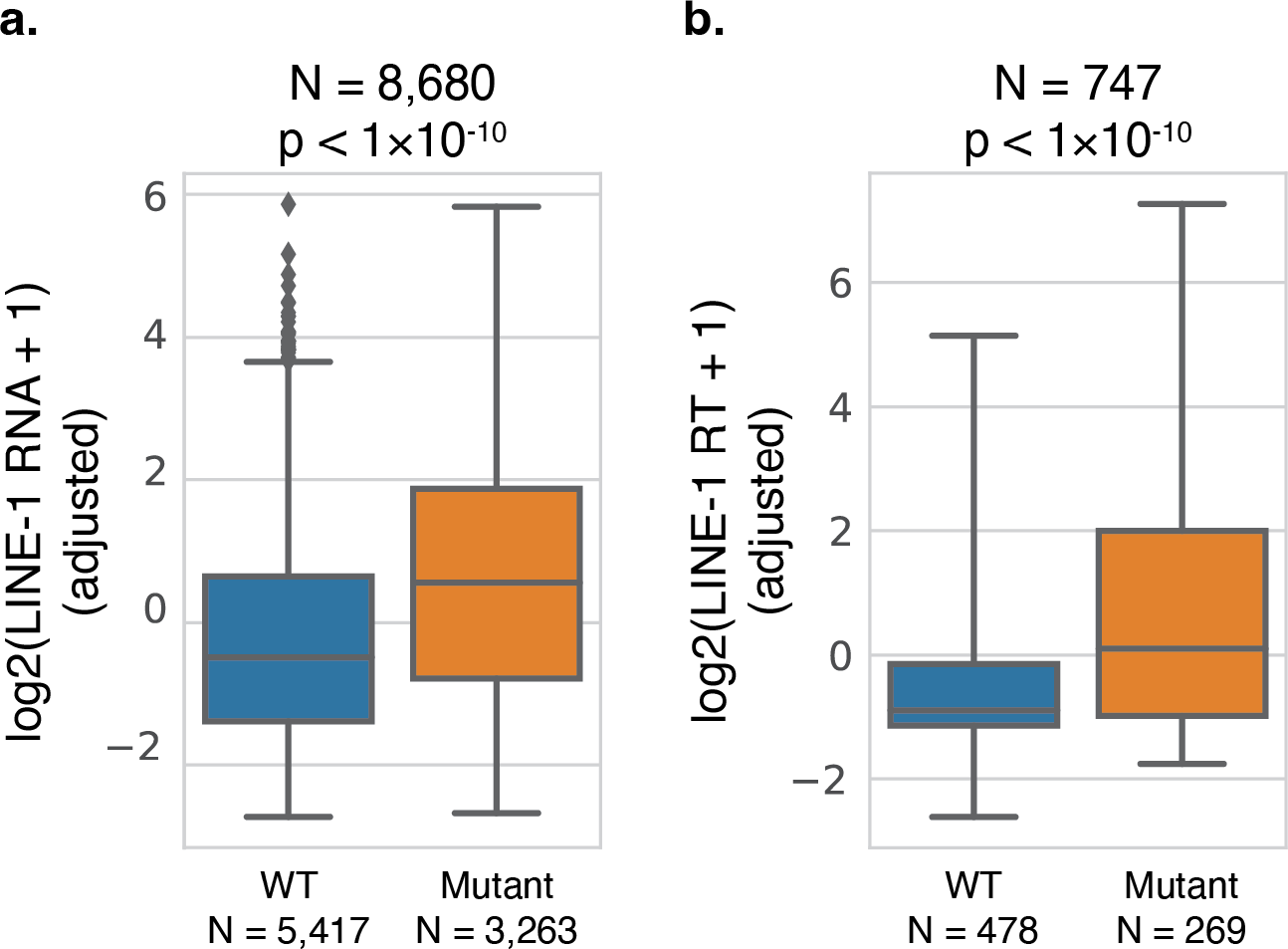
Stratification of L1 by p53 mutation. a) Expression of intact L1 RNA in samples with mutant or wild-type p53. Expression is adjusted for intronic rate as shown in Extended Data Fig. 3b. b) Count of somatic L1 retrotranspositions in samples with mutant or wild-type p53. Somatic RT counts are adjusted for sequencing covariates as shown in Extended Data Fig. 3c. a-b, Center line indicates median. Blue or orange box indicates interquartile range. Points more than 1.5 x IQR away from the IQR box are shown as individual outliers. P calculated from two-sided Mann- Whitney U test.

We previously created a “fitness model” of the selective advantage a *TP53* mutation gives to a tumor cell to quantify the pro-tumor advantage of loss of native p53 function conferred by missense mutations in a mathematical model (defined by the ability to bind 8 key transcriptional targets)^21^. The fitness model has been further extended here to represent the corresponding impact of additional types of alterations other than point mutations to *TP53* (see Methods) for the first time, enabling us to compute p53 functional fitness for every tumor sample in our dataset (Fig. 5a). We observe that L1 RNA expression and RT burden are each significantly correlated with mutant p53 fitness (R=0.30, p < 1x10^-10^; R=0.25, p < 1x10^-10^, respectively, Pearson correlation, Extended Data Fig. 4). Additionally, the correlation between L1 RNA and mutant p53 fitness remains significant when using L1 RT burden as a covariate, and the correlation between L1 RT burden and fitness similarly remains significant when using L1 RNA as a covariate.

**Figure 5.**
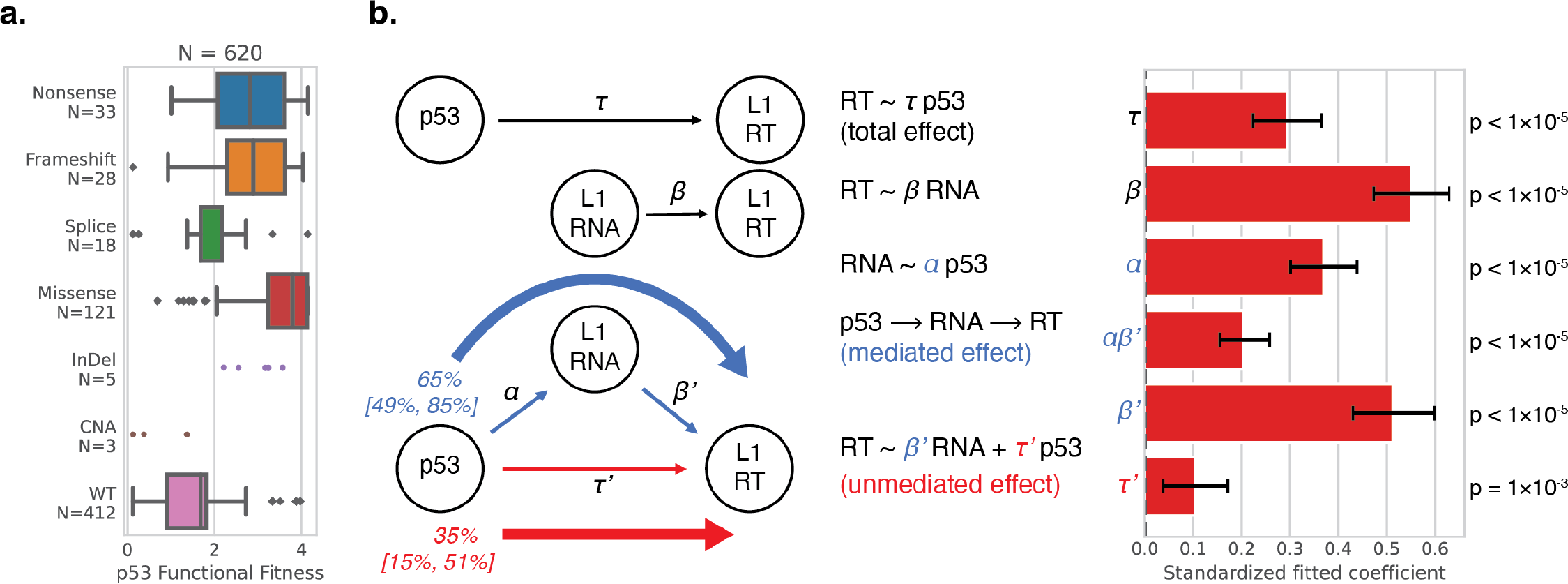
Mediation model relating p53 mutation, intact L1 RNA expression, and somatic L1 retrotransposition. a) p53 functional fitness score per sample by mutation category. InDel, in-frame insertion or deletion. CNA, copy number alteration. WT, wild type. Mutation categories with fewer than 10 samples are shown as individual points. Center line indicates median. Box indicates interquartile range. Points more than 1.5 x IQR away from the IQR box are shown as individual outliers. b) Mediation model taking p53 functional fitness as the independent variable, adjusted log2 of intact L1 RNA expression as the mediating variable, and adjusted log2 of somatic L1 retrotransposition count as the dependent variable. Left, schematic showing the linear regressions performed, as well as the resulting estimated weights for the mediated and unmediated pathways. 95% confidence intervals (square brackets) calculated with bootstrap resampling. Blue arrows, components of the mediated pathway. Red arrows, components of the unmediated pathway. Right, standardized fitted values for each coefficient within the mediation model and corresponding likelihood. Error bars indicate 95% confidence intervals calculated with bootstrap resampling, N=100,000. Coefficient estimates for α, β, τ, β’, τ’ estimated from ordinary least squares.

We next tested whether L1 RNA expression mediates the relationship between p53 mutational fitness and L1 RT burden. Statistical mediation analysis (see Methods) was performed on the subset of tumor samples for which WGS, RNA-seq, and p53 mutation data were all available (N = 620 tumor samples). The results revealed a significant mediated effect of p53 mutational fitness on L1 RT burden through L1 RNA (αβ’= 0.20, p < 1x10^-5^, Fig. 5b). The unmediated effect of p53 mutational fitness on L1 RT burden was also significant (τ’ = 0.10, p = 1.2x10^-3^). The combined linear regression model of L1 RT burden as a function of L1 RNA and p53 functional fitness has a Pearson correlation coefficient R = 0.56 (p < 1x10^-10^, Extended Data Fig. 5). This shows a complementary partial mediating role of L1 RNA expression on the relationship between p53 mutational status and L1 RT burden, suggesting that p53 regulates somatic retrotransposition via at least two mechanisms: one that is L1 RNA-dependent, and one that is not. This modeling at scale is thus highly consistent with the proposed dual regulatory role of p53 in restraining L1 retrotransposition^27^, with the non-L1 RNA mechanism likely mediated through regulation of genomic instability-associated processes.

## Discussion

Our study substantiates with quantitative statistical evidence that a regulatory relationship exists between p53 and L1 throughout human cancers. We have employed a theoretical fitness model to evaluate the extent to which mutation has altered p53 function in each tumor context, but the significance of the statistical mediation model is robust to the method for quantifying mutational status. Both the mediated and unmediated modulation of L1 retrotransposition by p53 remain significant when the model is evaluated using a binary mutated category (i.e., in every sample, p53 is either altered (1) or not (0)) as defined by cBioPortal (Extended Data Fig. 6)^25, 26^. While there are limitations when using observational data, such as cancer types examined, causal assumptions of the directionality of mediation, and survivorship bias of cancer cells with mutated p53 under L1 overexpression^28^, our large pan-cancer study is well-powered and is consistent with previous studies using human cell lines as model systems^19, 29^. Our analysis suggests that p53 transrepression of L1 through direct binding of the genomic L1 5’ UTR is likely to be somatically active across tissues and significantly mutagenically disrupted in cancers.

Many studies have noted that the role of p53 as a general regulator of cell cycle control, apoptosis, and senescence is insufficient to explain the extent of tumor suppression by p53^30–32^. In a 2018 review, Tiwari et al. hypothesized that p53 may also be suppressing tumor formation through its role in directly restricting expression of transposons^27^. We theorize here that various cellular stress response pathways may be activated by p53 in response to retrotransposition, and in addition p53 regulates L1 via different mechanisms, likely by acting on the L1 promoter that controls L1 RNA expression. Using the mechanistic models of stimulus-dependent regulation and tonic regulation^27^ as a framework for interpretation, our results suggest that L1 modulation in cancer by p53 uses both modes of regulation and occurs at both the transcriptional and retrotransposition levels. Previous studies have demonstrated mechanisms by which p53 can directly transrepress L1, by binding its 5’ UTR RNA polymerase II promoter and facilitating the deposition of histone marks^19, 20^. Our finding of a significant correlation between p53 mutational fitness and L1 RNA expression across a dataset of clinical tumor samples is consistent with this transrepression being somatically active and relevant to cancers. Repression of the L1 lifecycle at the transcriptional stage will logically preclude retrotransposition, as is represented by the RNA-mediated path of our statistical model. The non-RNA-mediated effect of p53 mutation modulating L1 retrotransposition may be imposed by negative selection against reintegrations^28^ or their consequences for gene expression^33–35^ following stimulus-dependent regulation by wild-type p53 (e.g., cell cycle arrest), or via a still undescribed mechanism.

To our knowledge, this work includes the first pan-cancer analysis of the expression levels of active L1 mRNA. Of note, a previous study observed relationships between L1 expression and DNA damage and replication stress, suggesting that the oncogenicity of L1 derepression is in excess of what can be attributed to completed cycles of retrotransposition that disrupt gene expression^36^, and therefore L1 RNA may itself be oncogenic through some mechanisms. For example, abundance of L1 transcripts can play a role in heterochromatin erosion^37^. Additionally, the presence of cytosolic L1 RNA/cDNA hybrids resulting from reverse transcription could lead to an inflammation response that can alter the tumor microenvironment by activating the cGAS innate immune receptor, which in turn activates the interferon pathway^38^. This analysis uncovers tissue- and tumor-type dependence of L1 expression at the RNA level and informs understanding of L1 behavior and activity in these different contexts. For instance, a previous study investigating L1 ORF1p binding in prostate cancer suggested the possibility that cytoplasmic ORF1p may affect RNA processing after finding many non-L1 transcripts associated with ORF1p, including in particular RNAs that were also enriched at p-bodies^38^. Although that study focused on prostate cancer, our analysis did not find significant activity of L1 retrotransposition in prostate tumors (only one somatic insertion was identified out of 15 tumor samples profiled), nor did intact L1 RNA expression in prostate tissues vary significantly between tumor and normal samples.

There are still many open questions about the interplay of L1 and p53 in cancer. In an earlier study (on a smaller cancer dataset), L1 ORF1p positively correlated with copy number alteration in breast, ovarian, and endometrial tumors, especially when p53 was mutated^36^. Our observations provide additional evidence into the diverse regulatory roles of p53 that influence L1 RT burden in cancer. Using state-of-the-art multi-modal modeling of L1 retrotranspositions (Total ReCall), L1 RNA expression (L1EM), as well as p53 mutational fitness, we are able to leverage statistical modeling of causality to uncover associations that were previously masked by experimental and biological heterogeneity. Having robust tools to measure L1 RNA expression, L1 retrotransposition, and p53 mutational fitness may prove useful in cancer risk stratification and advance personalized cancer medicine.

## Methods

### Pan-cancer datasets

#### TCGA

Whole genome sequencing data were downloaded from the Genomics Data Commons (GDC) cloud storage, using the GA4GH standard Data Repository Service (DRS) for URI resolution. DRS links to the hg19-aligned BAM files were collated from pre-existing Terra workspaces (https://app.terra.bio) for TCGA (v1.0 of ControlledAccess data). RNA-Seq data were downloaded from the GDC directly (https://portal.gdc.cancer.gov/repository) as hg38-aligned BAM files using the GDC Data Transfer Tool (https://docs.gdc.cancer.gov/Data_Transfer_Tool). All subsequently described processing of these data was performed within the Terra.bio cloud data platform (https://terra.bio/).

#### Genome in a Bottle

Alignments to hg38 and indices (BAM and BAI files, respectively) for PacBio and Illumina reads for the Genome in a Bottle project^22^ were downloaded from the NCBI archive. A complete list of access links can be found in Supplementary Table 1.

### Validation against long-read sequencing of LINE-1 retrotransposition

Non-reference L1 elements were called in the long read data using PALMER v 2.0.0^39^, and high confidence autosomal calls were selected. We then manually curated the calls by visually reviewing the PacBio reads at the loci of PALMER calls in IGV and identifying likely false negatives (non-reference L1 elements not called by PALMER in some of the members of the trio) and manually adding the calls in additional trio members according to the manual assessment (based on reads with insertions and/or clipped reads). L1 elements that were present in the genome of all the members of the son/father/mother trio were removed from consideration. We were left with 70 L1 elements present in the genomes of only one or two members of the trio of Ashkenazi Jewish ancestry (HG002/HG003/HG004) and 123 L1 elements present in the genomes of only one or two members of the trio of Han Chinese ancestry (HG005/HG006/HG007). As a sanity check of this curated call set, we verified that there were no L1 elements present in the genome of a child (HG002 or HG005) and absent from the genomes of both parents. This curated set of L1 elements was used for comparison with our L1 calls using Total ReCall on the short reads.

We ran Total ReCall (our Illumina short read-based L1 insertion caller described below) using different members of each trio as “case” and “control” (resulting in all 6 pairwise comparisons for each trio) and compared the results to the curated long-read calls as “ground truth”. For a “perfect” agreement between a PacBio and an Illumina call, we require that the call is present in all pairwise comparisons where it should be present with the correct filtering status (e. g., if a given L1 element is present only in the maternal genome in PacBio calls, it should be called in the mother vs. father and the mother vs. child and not called in any other of the 6 pairwise comparisons; if a given L1 element is present in the paternal and the child genomes in PacBio calls, it should be called in the father vs. mother and the child vs. mother, but also called in the father vs. child and the child vs. father yet filtered-out due to its presence in the “case” sample). In addition, we also assessed if the strandedness and the presence or absence of a L1 inversion agreed between the PacBIo and Illumina calls.

Overall, 137 out of 193 long-read calls were also called properly by Total ReCall, including 93 canonical retrotranspositions and 44 inversion-containing retrotranspositions. We inferred the transposon length for each of these 93 calls using PALMER calls as the distance between the middle points of the inferred confidence intervals for the start and end points in the L1 (thus excluding possible 3’ transductions) and compared it to the length inferred using Total ReCall. Agreement was very good (Pearson correlation R > 0.99, p < 1 × 10^-10^, t-test, Extended Data Fig. 1).

### Short-read whole-genome sequencing data from TCGA

#### Reprocessing of alignment files

Whole-genome sequencing reads data from The Cancer Genome Atlas (TCGA) were reverted to unaligned BAM format using the GATK v4.1.8.1 tool RevertSam with options “--SORT_ORDER queryname --VALIDATION_STRINGENCY SILENT”. Any reads that were not paired-end were dropped, and the remaining reads were converted to FASTQ format with GATK v4.1.8.1 tool SamToFastq with options “INTERLEAVE=true INCLUDE_NON_PF_READS=true”. The paired-end FASTQ files were then aligned to the hg38 human reference using BWA-MEM v0.7.15- r1140^40^ with options “-K 100000000 -p -v 3 -t 16 -Y -T 0”, where we used the “-Y” option to retain clipped read sequences for supplementary alignments that are clipped (i.e., “soft-clipping”), since clipped read information is the key evidence that Total ReCall leverages (see below). Duplicate reads were marked with GATK v4.1.8.1 tool MarkDuplicates using options “-- OPTICAL_DUPLICATE_PIXEL_DISTANCE 2500 --VALIDATION_STRINGENCY SILENT”.

#### Quality control data filtering

Variability of sequencing library quality across TCGA WGS data (which was generated over the span of years) can obscure the differentiable signal of biological relationships. With that in mind, we filtered the entire WGS dataset (4,314 samples from 2,106 patients) to only those samples sequenced with paired-end reads, where both mates had been retained in the alignment files stored in NCI Genomic Data Commons (4,142 samples from 2,058 patients). Each of these samples was realigned to the hg38 reference genome as described above. To de-duplicate individual patients with multiple tumor or normal samples, only the primary tumor and a single normal sample were retained, resulting in 1,848 tumor-normal pairs from 1,848 patients. Using all 1,848 tumor-normal pairs agnostic to additional data quality parameters, no correlation could be found between L1 RNA expression, p53 mutational status, and L1 retrotransposition, due to very large confounding differences between the libraries. To account for these differences and better capture biological relationships, the quality of each WGS library was evaluated using a modified Picard^41^ tool to quantify the total number of reads, average base quality, average read length, and fractions of reads with split or discordant alignments (the two sources of information on which Total ReCall relies). All reads that did not have a matched pair, that were marked as duplicates, that had mapping quality equal to zero, or whose base and quality score strings were inconsistent (i.e., of differing lengths) were removed from the analysis. From the remaining reads, for each sample, we calculated 1) the fraction of chimeric reads, 2) the fraction of overall clipped bases, 3) the average read length, 4) the average base quality, and 5) the total coverage. Note that a chimeric read is defined per Picard/GATK v4.1.8.1 as a read pair aligned in an unexpected orientation or significantly further apart than expected (with maximum insert size set to 100,000). Because Total ReCall will depend on signal from split-read alignments (i.e., clipped reads) to nominate candidate insertion sites, and because bwa mem is intended for use with reads a minimum of 70bp, tumor-normal pairs where either sample was sequenced with reads shorter than 70bp were removed from our analysis dataset. We additionally removed all paired samples with a chimeric read fraction in the tumor or normal sample greater than 2%, resulting in a total of 765 tumor-normal pairs from unique patients across 22 tumor types in our final analysis dataset.

Due to their importance, the five metrics for each tumor and paired normal sample calculated here were also used to adjust the retrotransposition count estimates for each tumor sample, as shown in Extended Data Fig. 3 and described below.

#### Identification of somatic LINE-1 retrotransposition

Our Total ReCall method for identification of somatic L1 insertions uses both soft-clipped reads, as well asreads not properly mapped as a pair (“discordant” read pairs) as a signal. We use the clipped reads as the primary signal because of their higher specificity: the last mapping point as well as the clipped sequence are the same (up to sequencing errors) for all the clipped reads supporting the same breakpoint. Interpretation of these types of signals arises from the mechanistic biochemistry of the life cycle of retrotransposons.

Class I transposable elements replicate via a reverse transcribed RNA intermediate (“copy-and-paste”) in a process known as retrotransposition, whereas class II elements replicate via a DNA intermediate (“cut-and-paste”). Further subdividing based on distinct mechanisms of mobility, there are three major categories of transposable elements: class I LTR retrotransposons (including endogenous retroviruses), class I non-LTR retrotransposons, and class II DNA transposons^42^. Among the transposable elements, only some non-LTR retrotransposons are known to be able to move within the genome. L1 elements are capable of autonomous retrotransposition, and SINE (e.g., Alu) and SVA elements are “parasites” that rely on the L1 machinery for mobility.

The difference between the mechanisms utilized by LTR and non-LTR retrotransposons is that the LTR retrotransposons first synthesize the double stranded DNA from RNA transcript in a complex process involving the long terminal repeats (LTRs) and then integrate that sequence into the host genome. On the other hand, enzymes of non-LTR retrotransposons (LINE elements) reverse transcribe their RNA directly into the genome using the so-called target-primed reverse transcription (TPRT) process^43^.

Unlike the LTR retrotransposons, which always integrate the complete provirus with the LTRs into the genome (sometimes, the internal provirus part may be removed by homologous recombination leaving a solo LTR), reverse transcription of a non-LTR retrotransposon may be terminated prematurely resulting in a partial integration. Apart from such a 5’ truncation, another common structural variant of L1 is inversion resulting from double priming and simultaneous reverse transcription of the same L1 transcript into both strands of the genome (Fig. 1d)^44, 45^. In addition to that, it is not uncommon for there to be 3’ transductions, where during L1 RNA expression the polymerase overruns the poly(A) signal at the 3’ end of the L1 element resulting in transcription and retrotransposition of some genomic sequence downstream of the 3’ end of the source L1 element in addition to the L1 element itself ^23, 46^. In the Total ReCall method developed here, we focus on detecting these three most common features of the structural variations that result from L1 retrotranspositions: 5’ truncation, L1 inversion, and 3’ transduction. We believe that the analysis of even rarer and more complex structural variations arising from L1 retrotransposons (e.g., instances with more than one template switching during the reverse transcription process, which would result in a series of inversions) is better left for long-read data.

We thus designed the Total ReCall algorithm (Fig. 1) to detect both “canonical” retrotransposition events and “inversion-containing” retrotranspositions. In particular, “single-stranded” retrotransposition (Fig. 1c) leads to a canonical retrotransposition with a single segment of L1 inserted into the genome (Fig. 1e), with the two ends of the insertion detectable in the clipped short-reads (Fig. 1g). On the other hand, a “double-stranded” retrotransposition event (Fig. 1d), where each strand of the target site genomic DNA incorporates reverse transcribed sequence of a single L1 mRNA, leads to an inversion-containing retrotransposition with a DNA insertion that contains part of a L1 sequence and another part of it that is inverted, i.e., contains reverse complementary L1 sequence, in an adjacent position in the genome (Fig. 1f), which will be reflected in the short-reads data (Fig. 1g).

To identify the breakpoints of the L1 insertion event, Total ReCall collects the clipped sequences of the reads in the vicinity, sort them by length in decreasing order and cluster in a cd-hit-like^47, 48^ way: take each new sequence and align it against the existing representatives; if a match is found, assign the sequence to the corresponding cluster, otherwise designate it as a new representative. We group the reads representing the “left” (clipped on the 3’ end w.r.t. the reference genome) and the “right” (clipped on the 5’ end w.r.t. the reference genome) breakpoints. The longest clipped sequence in the cluster serves as the representative of the breakpoint. If we orient the clipped sequence so that the clipping point is at its 5’ end, a canonical L1 retrotransposition is characterized by one breakpoint with the poly(T) sequence (regardless of whether a 3’ transduction is present) and the other breakpoint with a sequence mapping to the positive strand of the L1 sequence. The coordinate of the alignment of the clipped sequence at the breakpoint to the L1 sequence can be used to infer the length of the transposon (without the 3’ transduction, if one exists). An inversion-containing L1 retrotransposition is characterized by one breakpoint bearing the poly(T) sequence (regardless of whether a 3’ transduction is present) and the other breakpoint with a sequence mapping to the negative strand of the L1 sequence.

As noted above, we increased sensitivity by using the “soft clipping” instead of the “hard clipping” for supplementary alignments (*bwa* option *-Y*) relevant when the longer part of the read spanning the breakpoint contains the L1 sequence (and whose primary mapping is thus to L1 genomic sequence elsewhere in the genome). The use of such supplementary alignments is highly desirable since they provide a longer clipped sequence, which can in turn be aligned to the L1 sequence more specifically.

Total ReCall also identifies reads that belong to discordant read pairs and map near the breakpoints and use their mates as a secondary signal to determine the confidence of the call. If such a read is also clipped at the breakpoint location, its mate is given a higher preference over the mates of the other reads.

Clipped sequences, as well as sequences of the reads belonging to discordant read pairs, were aligned to the L1 consensus sequence using LAST aligner^49^. The consensus sequence of the L1 transposon was constructed by merging the three DFAM^50^ sequences for its 3’, 5’ ends and ORF2 subdomain (DF0000225, DF0000226, DF0000316). We resolved the ambiguous characters in the DFAM consensus sequences using the alignment of the 146 intact L1 sequences from L1base^24^.

To identify somatic L1 retrotranspositions (i.e., in our typical use case of tumor-specific insertions), Total ReCall checked each call in the case sample for the presence in its control sample of the signal of a clipped read with a matching sequence or a breakpoint, retaining as “somatic” (i.e., case-specific) only calls without such signal in the control. In addition, we implemented filters for low complexity and regions with large numbers of alignment artifacts (“high entropy regions”), e.g., regions having too many clipped reads with distinct sequence and low complexity regions where genomic sequence matches clipped sequence. Finally, Total ReCall requires each of the 2 breakpoints of a called insertion rely on at least 3 clipped reads.

It is worth mentioning that the short-read data possess inherent limitations, for example, difficulty identifying 3’ transductions. Long such transductions may be identified via discordant read pairs whose mates map downstream of source LINE-1 element. At the same time, this method will fail to identify short 3’ transductions. Some other shortcomings are the ambiguity of pairing between the left/right breakpoints if multiple ones are present in the same region as well as low mappability of some genomic regions (in particular, pericentromeric regions).

### RNA sequencing

#### Reprocessing of alignment files

Public RNA-seq data from TCGA was reverted to unaligned FASTQ format using GATK v4.1.8.1 tools RevertSam with options “--SORT_ORDER “queryname” --VALIDATION_STRINGENCY SILENT” and SamToFastq with options “INTERLEAVE=false INCLUDE_NON_PF_READS=true”. Any reads that were not paired were dropped. The paired-end FASTQ files were then aligned to the hg38 human reference genome using STAR v2.7.9a.

#### Quality control data filtering

Alignment files for a total of 10,904 RNA-seq samples from 10,089 individuals (10,174 tumor samples and 730 normal samples) were downloaded from GDC as described above. 28 of these (all tumor samples) could not be reverted to fastq for realignment due to the presence of unpaired reads or otherwise corrupted downloaded alignment files. The remaining 10,876 samples were realigned as described above. 634 of these (623 tumor samples and 11 normal samples) were sequenced with single-end reads, and therefore removed from our dataset. 59 additional tumor samples were removed from our dataset for containing a strand-specific sequencing library. Finally, we filtered out any metastatic, recurrent, or new primary tumor samples and deduplicated the patient samples in the dataset to include no more than one primary tumor and one normal sample per patient, resulting in a dataset of 9,011 tumor samples and 719 normal samples from 9,081 individuals across 32 tumor types.

#### LINE-1 RNA quantification

L1 RNA expression was quantified in the 9,011 tumor and 719 normal RNA-seq samples using L1EM^24^, which formalizes a framework for quantification of expression that is based on the mechanisms of active transcription of L1 elements. Read counts for proper expression at loci corresponding to the L1Base v2 database^24^ of “FLI-L1s” (L1s that are full length and intact in both ORFs) were converted to transcripts per million (TPM) of active, fully-intact L1 expression.

#### Counting L1HS alignments

To compare performance of L1 RNA quantification by L1EM, we also estimated L1 expression using samtools view v1.3.1 with options “-F 1540” and “-L” to pass coordinates in bed format of L1HS annotations from RepeatMasker open-4.0.5 Repeat Library 20140131^3^.

From the output of alignments which overlapped the input L1HS coordinates, unique qnames were counted, and normalized against the total unique qnames in the alignment file to calculate an approximate expression level in reads per million (RPM) for every sample (Extended Data Fig. 7). The samples are noticeably more homogeneous than when using L1EM (Fig. 3a), and the largest increase in median tumor estimate over median normal estimate occurs in stomach adenocarcinoma, where the tumor median is only 13% higher than the normal median.

### Categorizing tumors as p53 wild type or mutant

Categorical designations of p53 alteration were obtained through the public repository cBioPortal (www.cbioportal.org, accessed 23 February 2023) by querying “TCGA PanCancer Atlas Studies” (which includes 10,967 samples from 10,953 patients in 32 studies) for all alterations in “TP53”. A sample will be marked as “altered” in TP53 if any non-synonymous mutations, amplification, deep deletion, or structural variants have been identified in that sample. Shallow deletions or low-level copy number gains of the gene do not contribute to classification of a sample as altered. The altered annotations were used to divide tumor samples into p53 mutant and wild type categories to test significance with Mann-Whitney U tests as shown in Fig. 4, and to demonstrate robustness of the mediation model between p53, L1 RNA, and L1 RT burden as shown in Extended Data Fig. 6.

### Adjusting RNA and RT estimates based on sequencing metrics

Intronic rate of RNA-seq, which has previously been shown to be a key confound for L1 expression^51^, was calculated using RNA-SeQC v2^52^. A linear regression between the L1 RNA estimates and the intronic rate was then performed in Python v3.7.12 using statsmodels v0.13.0 OLS to perform an ordinary least squares regression. The residuals from this model were then used as the adjusted L1 RNA estimates (Extended Data Fig. 3b).

Similarly, quality metrics for the WGS samples were calculated as described above. An ordinary least squares model was fitted to the tumor-specific retrotransposition counts as a function of the average read length, depth of coverage, average base quality, chimeric read fraction, and clipped base fraction in both the tumor sample and the paired normal sample. The residuals from this model were then used as the adjusted L1 RT estimates (Extended Data Fig. 3c).

### Quantifying the functional fitness of p53

Somatic *TP53* mutations, copy number variation data, and the variant allele fractions for *TP53* mutations in TCGA samples were downloaded from the National Cancer Institute’s Genomic Data Commons repository^53^. The variant allele fractions were averaged across mutation callers. Functional fitness of missense mutations was scored as defined in previous work^21^. Reverse-phase protein assay data and copy number alteration data were utilized to infer mutant- and tissue-specific p53 protein concentrations as previously described^21^.

We determine the contribution of p53 to tumor fitness as follows. Wild-type p53 acts as a transcription factor and regulates DNA damage by transactivating pro-senescence/apoptotic genes. Mutations in p53 often result in loss of DNA-binding ability to varying degrees. We estimate the oncogenic advantage of having a p53 mutation by the probability of mutant p53 not binding eight principal transcriptional targets (*WAF1*, *MDM2*, *BAX*, *h1433s*, *AIP1*, *GADD45*, *NOXA*, and *P53R2*). The probability of not binding target DNA depends on the concentration of the protein and the affinity of the mutation to the target site. Importantly, the concentration of p53 depends on the tissue. In this work, we utilize the method described in Hoyos et al. 2022 for missense mutations^21^. For wild-type p53, we compute the fitness in an analogous fashion to the missense mutations, except that we use wild-type p53 concentrations and wild-type p53 affinities to DNA promoter sites. We extend this method to other mutations as follows. For silent/non-coding mutations, we consider the fitness to be wild type. For nonsense and frameshift mutations, we consider the lack of binding to be dependent on the position of the first altered residue along the protein. Specifically, we compute the fitness for these mutations as follows:

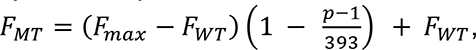

where *F*_*MT*_ is the fitness of the mutation, *F*_*WT*_ is the fitness of wild-type p53, *F*_*max*_ is the maximum fitness (which corresponds to a complete deletion of the p53 gene), and *p* is the position of the mutation (which may take an integer value between 1 and 393, inclusive). In this way, a position of “1” corresponds to maximum fitness, and a position of “394” (which would be after the protein) corresponds to wild type fitness. Tumors with wild-type p53 are assigned the wild-type p53 fitness that corresponds to the tissue in question.

### Mediation model

Statistical analysis was performed in Python v3.7.12 using statsmodels v0.13.0 OLS to evaluate the mediation model. For the 620 samples for which we had WGS, RNA-seq, and p53 mutation data available, five linear regressions were fitted (see Fig. 5b):

1. adjusted log2(RT) = τ x p53 mutation score + c_1_
2. adjusted log2(RT) = β x adjusted log2(RNA) + c_2_
3. adjusted log2(RNA) = α x p53 mutation score + c_3_
4. adjusted log2(RT) = β’ x adjusted log2(RNA) + τ’ x p53 mutation score + c_4_
5. adjusted log2(RNA) = β* x adjusted log2(RT) + α* x p53 mutation score + c_5_

Note that only equations (3) and (4) are necessary for quantifying the magnitude and significance of the mediation. Equation (1) is used to normalize the mediated and unmediated effects to the total effect of p53 on L1 RT burden. Equation (2) is included here to evaluate the correlation between L1 RNA and RT burden alone, without considering p53. Equation (5) (results not shown in Fig. 5 for simplicity) was evaluated to confirm that p53 has a significant effect on L1 RNA even when controlling for L1 RT burden. In each model, c_i_ incorporates both the intercept and the error terms. The fitted coefficient values were then standardized based on the estimated standard deviations for each variable^54^. The ratio of τ’ to τ gives the estimate for the percentage of the total impact of p53 on L1 RT burden that is not mediated by L1 RNA. The product of coefficients α and β’ (which is equivalent to τ – τ’) gives a coefficient for the effect of p53 on L1 RT burden via L1 RNA, and similarly the ratio of αβ’ to τ gives the estimate for the percentage of the total impact of p53 on L1 RT burden that is mediated by L1 RNA. 95% confidence intervals for all estimates were calculated using bootstrap resampling of the samples, N=100,000.

## Code availability

Total ReCall code for LINE-1 retrotransposition detection from short-read whole-genome sequencing data will be made publicly available on GitHub at the time of peer-reviewed publication.

## Acknowledgements

The results shown here are primarily based upon data generated by the TCGA Research Network: https://www.cancer.gov/tcga.

---

**Extended Data Figure 1.**
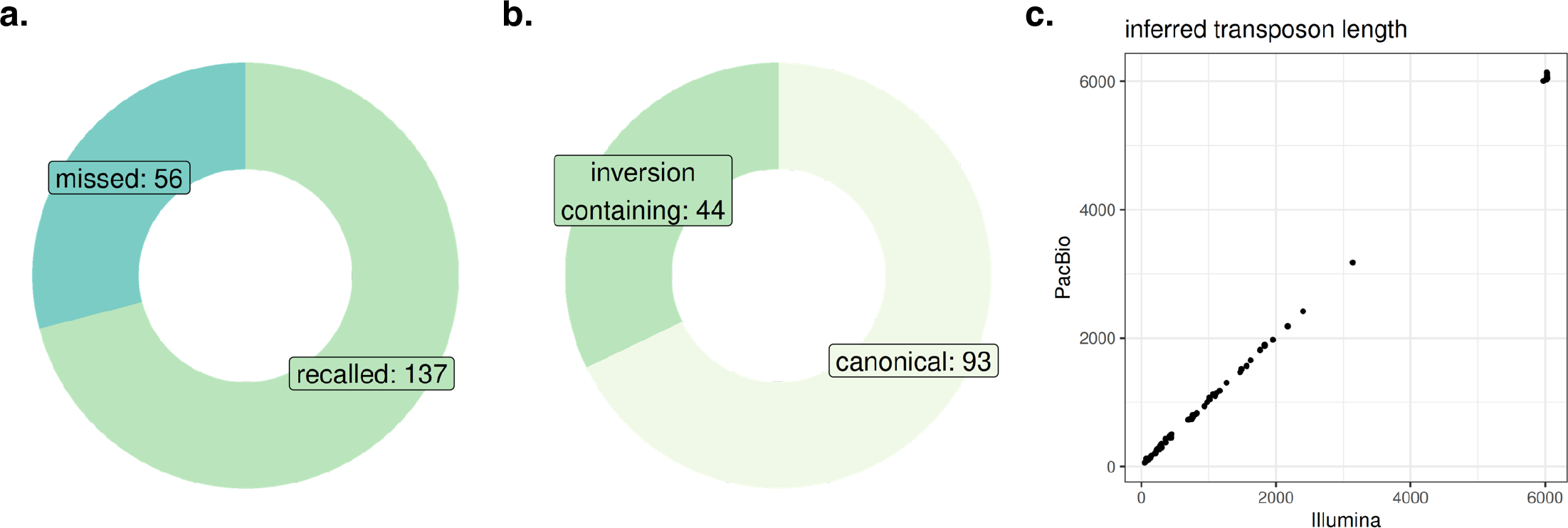
Genome in a Bottle validation of Total ReCall. a) Breakdown of curated L1 calls identified in PacBio reads by PALMER that are perfectly recalled by Total ReCall in Illumina short-read calls. N = 193 total curated calls. Green, 137 long-read calls identified by short reads. Blue, 56 long-read calls missed by short reads. b) Within the long-read calls identified by short reads, breakdown of canonical and inversion-containing insertions. c) Estimated length of canonical L1 insertions from short-read Illumina data vs. observed insertion length in PacBio data. N = 93, R > 0.99, p < 1 x 10^-10^, Pearson correlation.

**Extended Data Figure 2.**
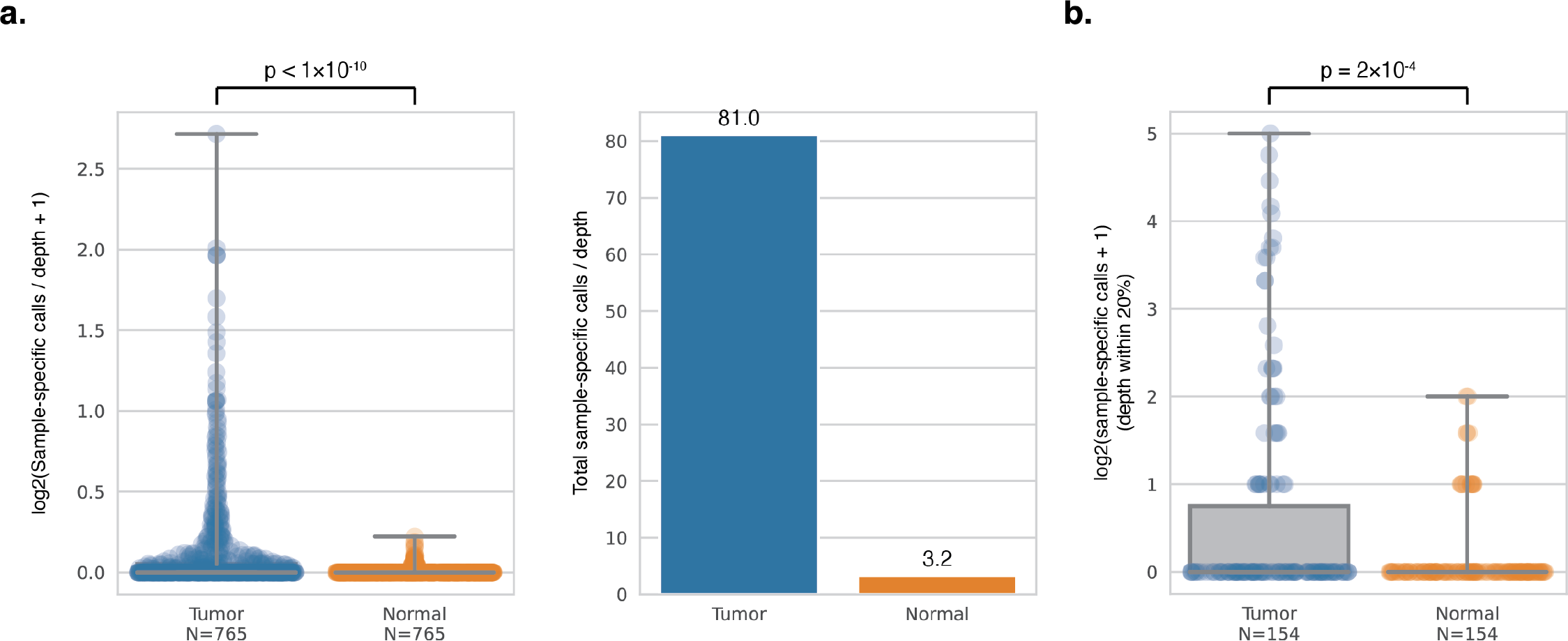
Adjusting tumor- and normal-specific calls per sample by sequencing depth. a) Left, ratio-normalizing the calls in each sample by the average read depth in that sample. Right, total of all calls per average read depth for all 765 tumor or normal samples. Y-axes in units of number of calls per read depth. N = 765 tumor and normal samples. b) Actual counts of calls from subset of dataset where tumor and normal sample were sequenced to comparable depth (within 20%). N = 154 tumor and normal samples. a-b) Center lines indicate median. Gray boxes indicate interquartile range. Whiskers extend to minimum and maximum values. Individual points shown for every sample. P calculated from one-sided Mann-Whitney U test. Blue, tumor samples. Orange, normal samples.

**Extended Data Figure 3.**
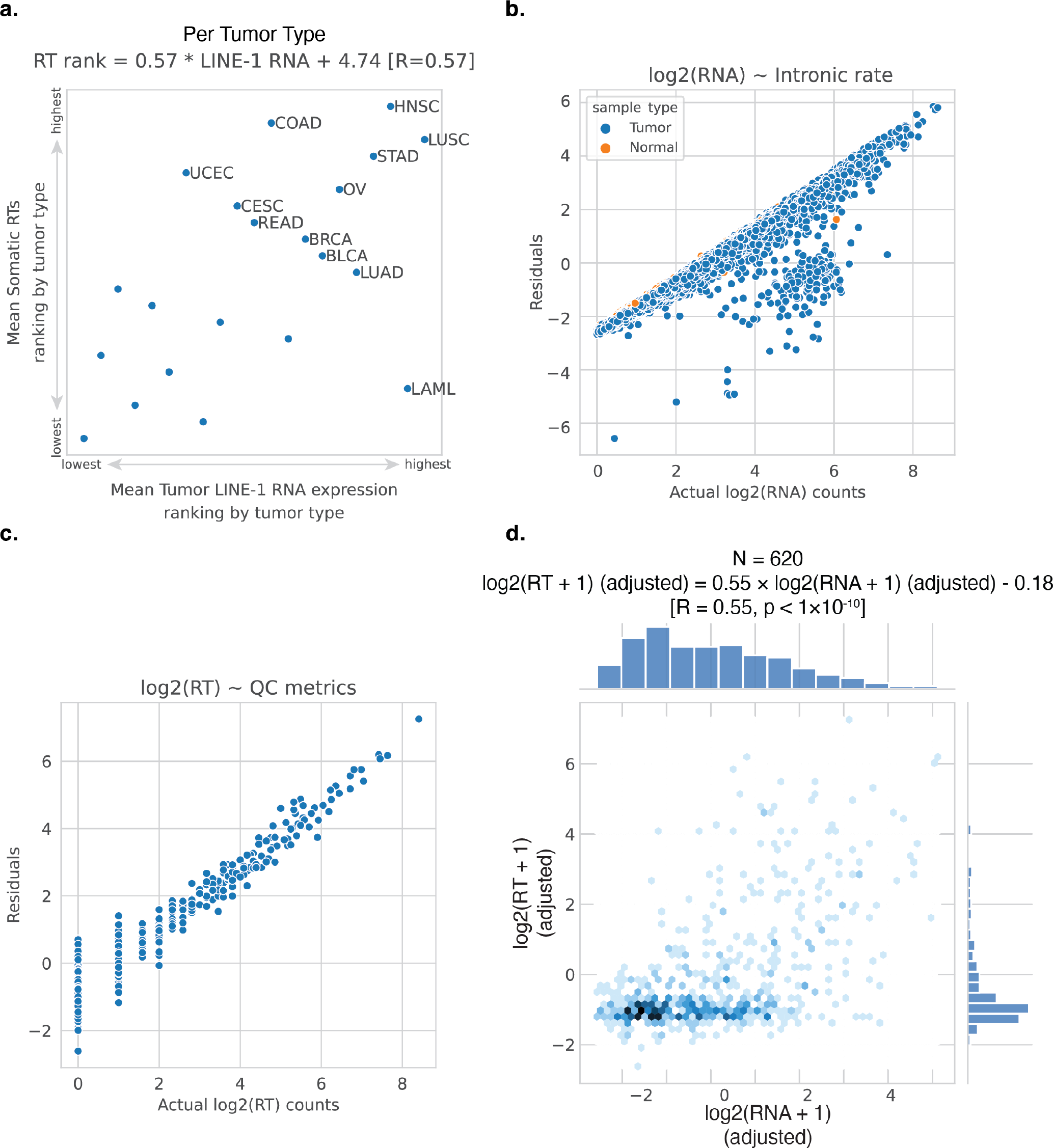
Relating L1 RNA expression and L1 RT. a) Ranking tumor types based on highest to lowest average count of somatic RTs and highest to lowest average expression of intact L1 RNA. Points representing tumor types with an average of less than 1 RT per sample are not labeled, with the exception of lung adenocarcinoma (LUAD) and acute myeloid leukemia (LAML) due to their high rankings for L1 RNA expression. Labels correspond to TCGA study abbreviations. N = 21 tumor types. Pearson correlation coefficient R = 0.57. b-c) Adjusted measurements based on residuals from linear regression model vs. raw measurements. b) Adjusted estimates of intact L1 RNA expression per sample (log2 TPM) based on intronic rate of RNA-seq. N = 9,730 tumor and normal samples. c) Adjusted counts of somatic L1 RT per sample (log2 count) based on total coverage, average base quality, read length, rate of clipped bases, and rate of chimeric alignments in both the tumor and paired normal WGS. N = 765 tumor samples. d) Correlation between adjusted L1 RT and adjusted L1 RNA per tumor sample. N = 620 tumor samples, R = 0.55, p < 1 x 10^-10^, Pearson correlation.

**Extended Data Figure 4.**
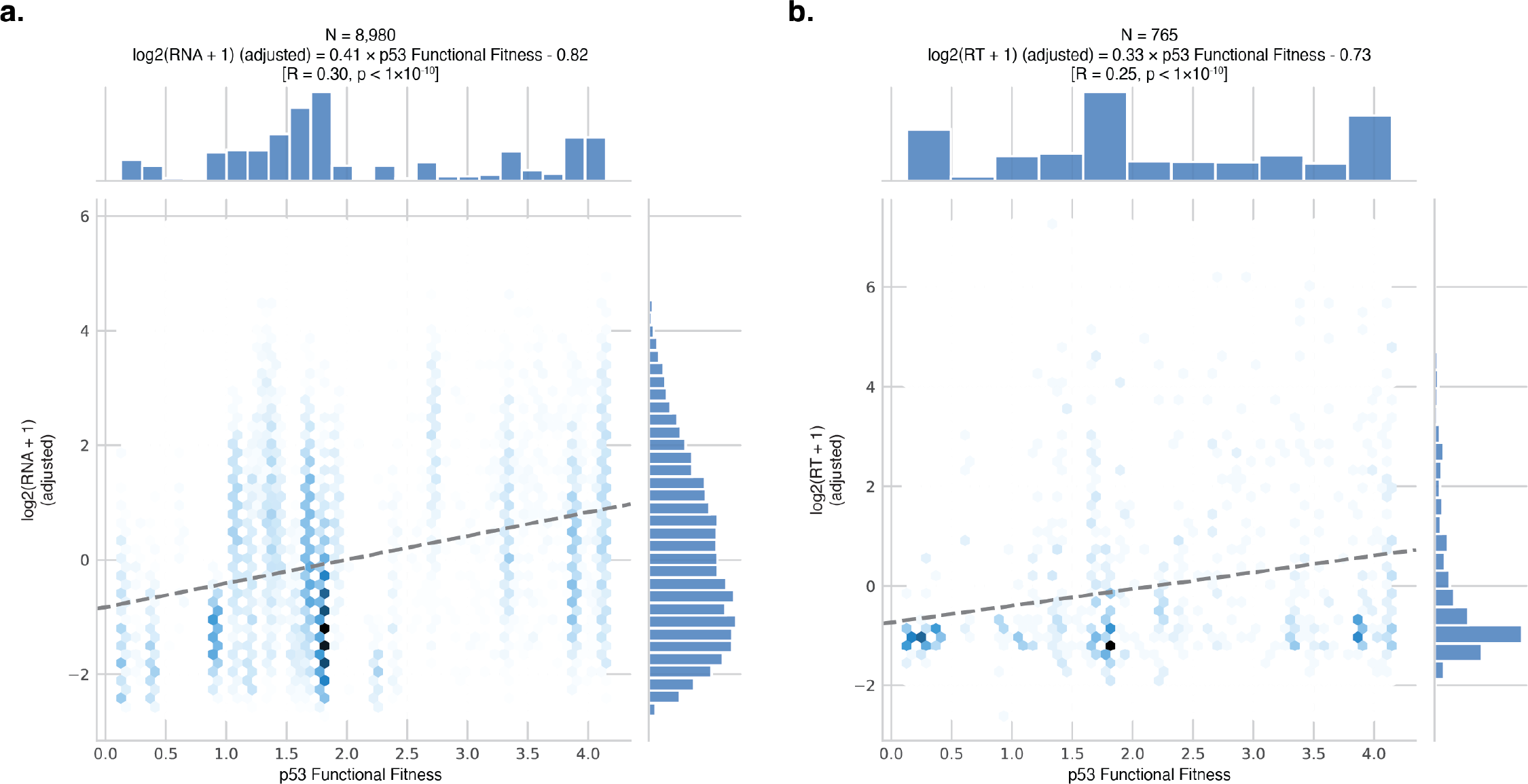
Correlation between p53 functional fitness and L1. a) Correlation between p53 functional fitness and expression of intact L1 RNA (log2 TPM, adjusted). All tumor samples with RNA-seq and available *TP53* mutation information are included; N = 8,980. R = 0.30, p < 1 x 10^-10^, Pearson correlation. b) Correlation between p53 functional fitness and count of somatic L1 retrotranspositions (log2 count, adjusted). All tumor samples with WGS and available *TP53* mutation information are included; N = 765. R = 0.25, p < 1 x 10^-10^, Pearson correlation. a-b) Dashed lines show fitted linear regression model, ordinary least squares.

**Extended Data Figure 5.**
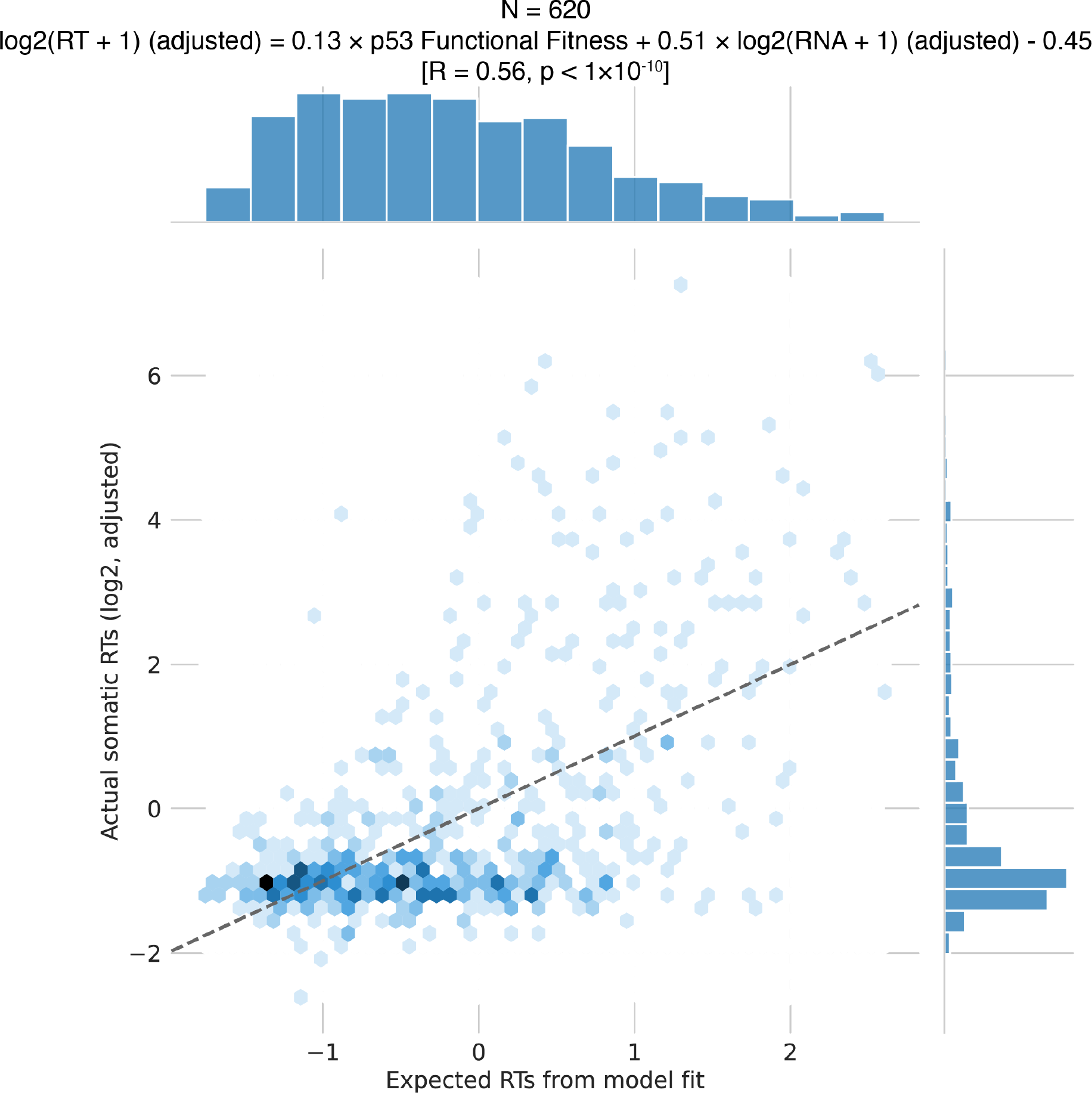
Correlation between L1 RT and mediation model fit. N = 620, R = 0.56, p < 1 x 10^-10^, Pearson correlation. Dashed line, y=x.

**Extended Data Figure 6.**
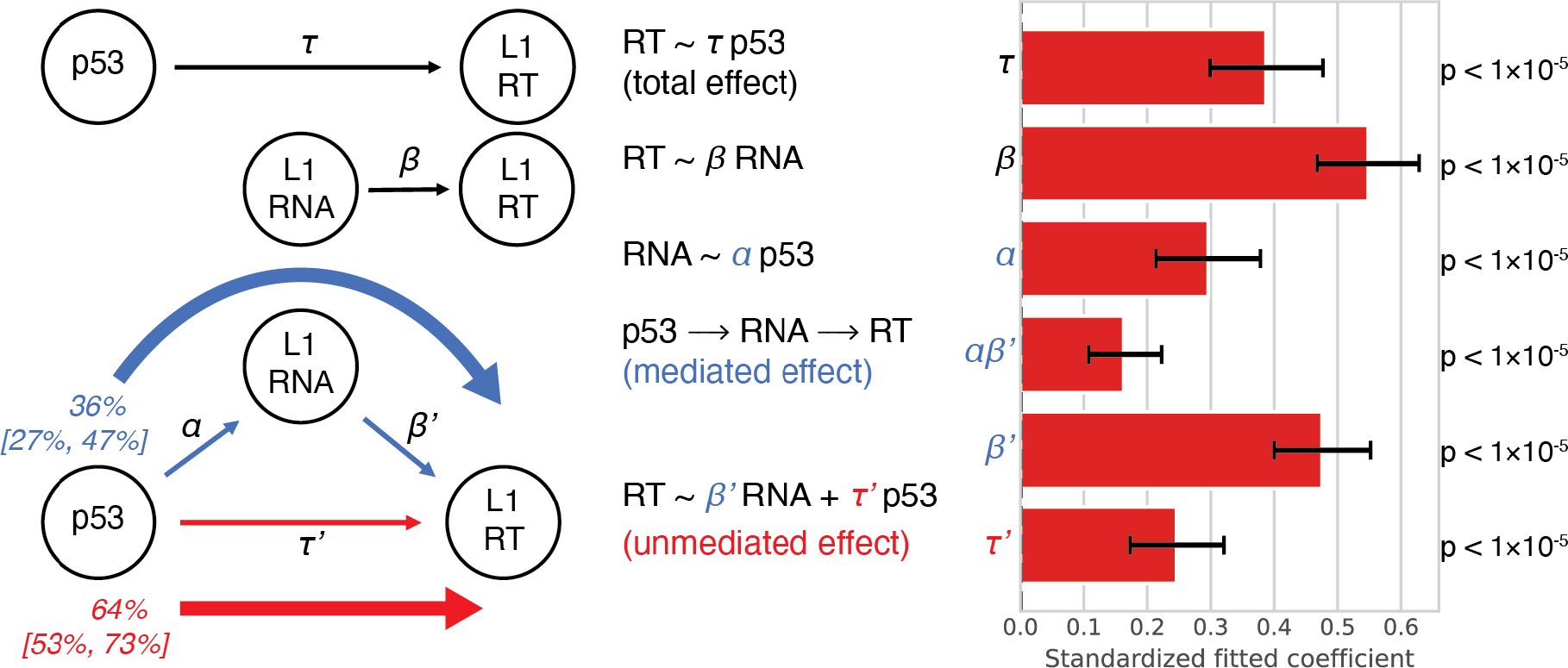
Mediation model taking *TP53* alteration status as the independent variable. In contrast to Fig. 5 (which uses p53 functional fitness), adjusted log2 of intact L1 RNA expression as the mediating variable, and adjusted log2 of somatic L1 retrotransposition count as the dependent variable. Schematics and labels as in Fig. 5b.

**Extended Data Figure 7.**
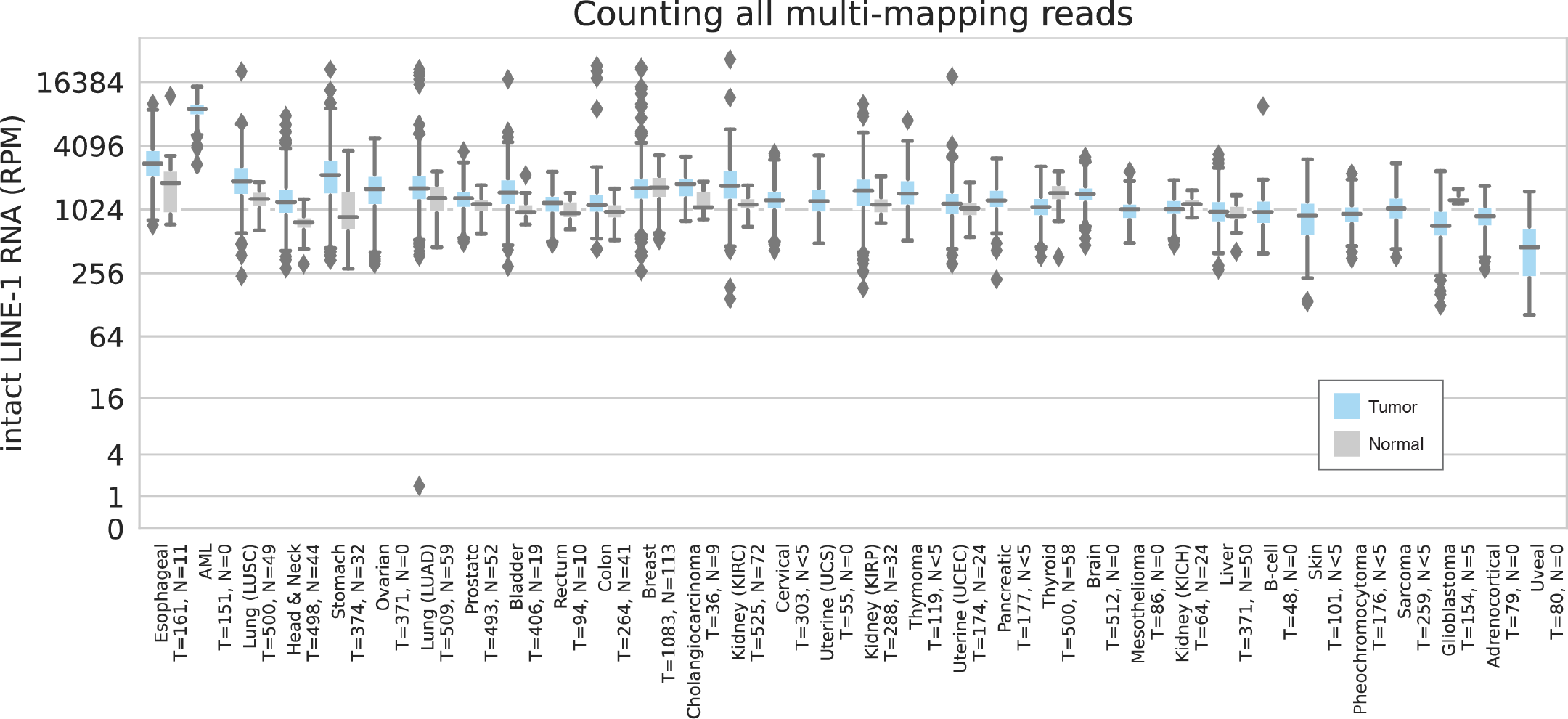
Estimating intact L1 RNA expression from read overlaps with RepeatMasker L1HS annotations in each sample by tumor type. Sample counts, plots, and tissue type ordering are as in Fig. 3a.

